# The N-terminal domain of spike glycoprotein mediates SARS-CoV-2 infection by associating with L-SIGN and DC-SIGN

**DOI:** 10.1101/2020.11.05.369264

**Authors:** Wai Tuck Soh, Yafei Liu, Emi E. Nakayama, Chikako Ono, Shiho Torii, Hironori Nakagami, Yoshiharu Matsuura, Tatsuo Shioda, Hisashi Arase

## Abstract

The widespread occurrence of SARS-CoV-2 has had a profound effect on society and a vaccine is currently being developed. Angiotensin-converting enzyme 2 (ACE2) is the primary host cell receptor that interacts with the receptor-binding domain (RBD) of the SARS-CoV-2 spike protein. Although pneumonia is the main symptom in severe cases of SARS-CoV-2 infection, the expression levels of ACE2 in the lung is low, suggesting the presence of another receptor for the spike protein. In order to identify the additional receptors for the spike protein, we screened a receptor for the SARS-CoV-2 spike protein from the lung cDNA library. We cloned L-SIGN as a specific receptor for the N-terminal domain (NTD) of the SARS-CoV-2 spike protein. The RBD of the spike protein did not bind to L-SIGN. In addition, not only L-SIGN but also DC-SIGN, a closely related C-type lectin receptor to L-SIGN, bound to the NTD of the SARS-CoV-2 spike protein. Importantly, cells expressing L-SIGN and DC-SIGN were both infected by SARS-CoV-2. Furthermore, L-SIGN and DC-SIGN induced membrane fusion by associating with the SARS-CoV-2 spike protein. Serum antibodies from infected patients and a patient-derived monoclonal antibody against NTD inhibited SARS-CoV-2 infection of L-SIGN or DC-SIGN expressing cells. Our results highlight the important role of NTD in SARS-CoV-2 dissemination through L-SIGN and DC-SIGN and the significance of having anti-NTD neutralizing antibodies in antibody-based therapeutics.

## Introduction

Coronavirus disease 2019 (COVID-19) is caused by the novel human coronavirus, a severe acute respiratory syndrome coronavirus 2 (SARS-CoV-2)^1,2^. SARS-CoV-2 is an enveloped virus and requires specific cellular receptors to infect cells in a manner similar to that observed in other enveloped viruses such as the influenza virus and herpesviruses^3–7^. The spike glycoprotein of SARS-CoV-2 associates with ACE2 on host cells and mediates membrane fusion with the host cell membrane during infection^3,4^. The spike glycoprotein is organized as a homotrimer and each chain is composed of two main segments, the S1 and S2 subunits. The S1 subunit is responsible for receptor binding, whereas the S2 subunit mediates membrane fusion^8,9^. Within the S1 subunit are two domains known as the N-terminal domain (NTD) and receptor-binding domain (RBD). The RBD is recognized as the main domain that interacts with the ACE2 receptor on the host cells and antibodies against RBD block infection^9^. Although the function of surface-exposed NTD that is structurally coupled with the RBD of the neighboring chain still remains unclear, certain human anti-NTD monoclonal antibodies neutralize SARS-CoV-2 infection^10^. This suggests that NTD as well as RBD play a pivotal role in SARS-CoV-2 infection.

Pneumonia is the primary complication associated with SARS-CoV-2 infection in severe cases^11^. Although ACE2 is highly expressed in the intestine and kidney, the expression of ACE2 in the lung is limited, and relatively low levels of ACE2 expression are observed only in type-II alveolar cells^12^. Therefore, it has remained unclear how the low levels of ACE2 in the lung are involved in pneumonia caused by SARS-CoV-2 infection. In addition, a single-cell RNA sequencing study of bronchoalveolar-lavage samples revealed that viral RNA could be detected in cell populations that did not express ACE2^13^. Taken together, this evidence indicates that ACE2 expression alone cannot explain the tropism of SARS-CoV-2 infection, and alternative entry receptors might be involved in SARS-CoV-2 infection of ACE2-low or deficient cells. Besides, SARS-CoV-2 is also detected in multiple non-pulmonary organs such as the intestine, liver, kidney, and blood vessels of COVID-19 patients^14^. Based on these observations, we sought to identify a potential alternative receptor for the SARS-CoV-2 spike protein using a lung cDNA library and discuss the mechanisms of SARS-CoV-2 dissemination within the host.

## Results

### Lung cDNA library screening identified L-SIGN as a receptor for NTD of the SARS-CoV-2 spike protein (SCoV2-NTD)

In order to identify a receptor in the lung for the SCoV2-spike protein, we used expression cloning with a lung retrovirus cDNA library. Cells transfected with a lung cDNA library were stained with SCoV2-NTD human IgG-Fc fusion protein (SCoV2-NTD-Fc) as well as RBD-Fc fusion protein (SCoV2-RBD-Fc) and the stained cells were enriched using a cell sorter (Fig. 1a). After two rounds of enrichment, 83.7% of the cells were stained with SCoV2-NTD-Fc (Fig. 1b) while we could not obtain cells stained with SCoV2-RBD-Fc even after three rounds of enrichment. We obtained single-cell clones that were stained with SCoV2-NTD-Fc and amplified the genes derived from the library using PCR. Notably, L-SIGN (CD299) was identified from all the clones that were stained with SCoV2-NTD-Fc (Fig. 1c). Following this observation, the cell clones were stained with antibodies against L-SIGN (Fig. 1d). These data suggested that L-SIGN is a major molecule that interacts with SCoV2-NTD in the lung. Although L-SIGN is mostly expressed in the liver, the lung has the second-highest expression of L-SIGN according to the RNA-seq tissue data obtained from the Genotype-Tissue Expression (GTE) project (Suppl. Fig. 1).

**Fig. 1.**
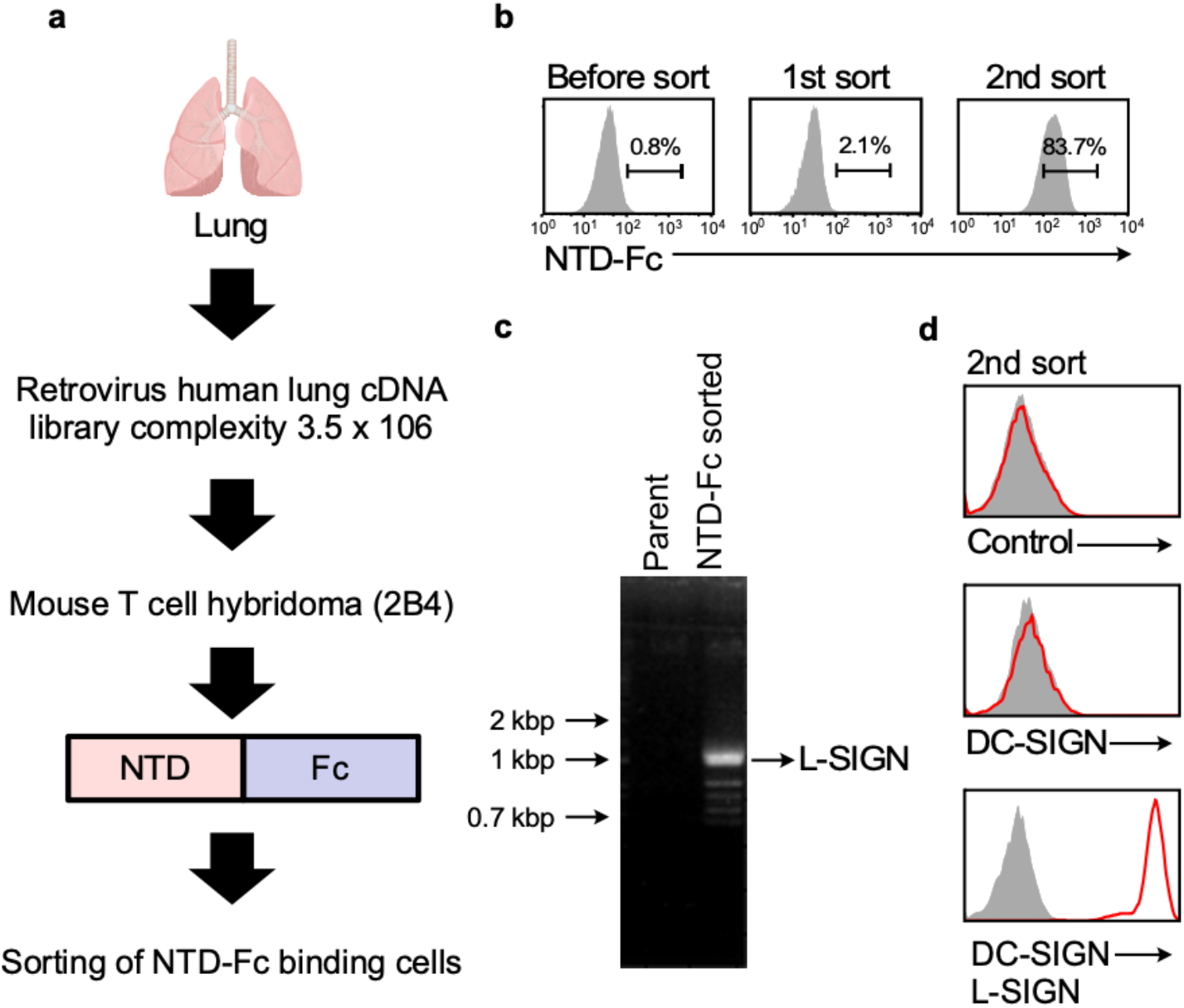
Lung cDNA library screening using the NTD-Fc of SCoV2 spike. **a,** The workflow of lung cDNA library screening. **b,** Cells transfected with retrovirus cDNA library were stained with SCoV2-NTD-Fc. Proportions of cells stained with SCoV2-NTD-Fc are shown. **c,** DNA agarose gel electrophoresis of cDNA library derived genes amplified from a single-cell clone stained with SCoV2-NTD-Fc. The indicated band was identified as L-SIGN. **d,** Single-cell clone stained with SCoV2-NTD-Fc was analyzed by anti-DC-SIGN mAb and anti-DC/L-SIGN mAb (red line) or control (shaded gray).

### NTD of the spike facilitates the binding of SCoV2 to L-SIGN and DC-SIGN via unique glycans

L-SIGN shares a high amino acid sequence homology and functional similarity with DC-SIGN that is also present in the lung (Suppl. Fig. 1). On the other hand, CD147 has been reported to be involved in SARS-CoV-2 infection^15^. Accordingly, we transfected L-SIGN, DC-SIGN, ACE2, and CD147 into HEK293T cells and evaluated the binding of NTD- or RBD-Fc fusion proteins (Fig. 2a). We observed that the NTD-Fc bound to both surface-expressed L-SIGN and DC-SIGN transfectants but not to ACE2 and CD147 transfectants. The RBD-Fc fusion protein bound to ACE2 but not L-SIGN, DC-SIGN, or CD147 transfectants. This indicates that the binding of NTD-Fc to L-SIGN or DC-SIGN is mediated by NTD and not the Fc portion. Anti-DC-SIGN mAb blocked the binding of NTD-Fc to DC-SIGN transfectants but not to L-SIGN transfectants (Fig. 2b). L-SIGN and DC-SIGN have been shown to bind to intercellular adhesion molecule 3 (ICAM-3) through sugar chain recognition. In addition, mannan is known to be a ligand of both L-SIGN and DC-SIGN. When DC- or L-SIGN transfectants were preincubated with mannan, the binding of NTD-Fc to DC-SIGN and L-SIGN was drastically reduced (Fig. 2b). These data indicate that unique glycans on NTD of SARS-CoV-2 are involved in the interaction with L-SIGN and DC-SIGN.

**Fig. 2.**
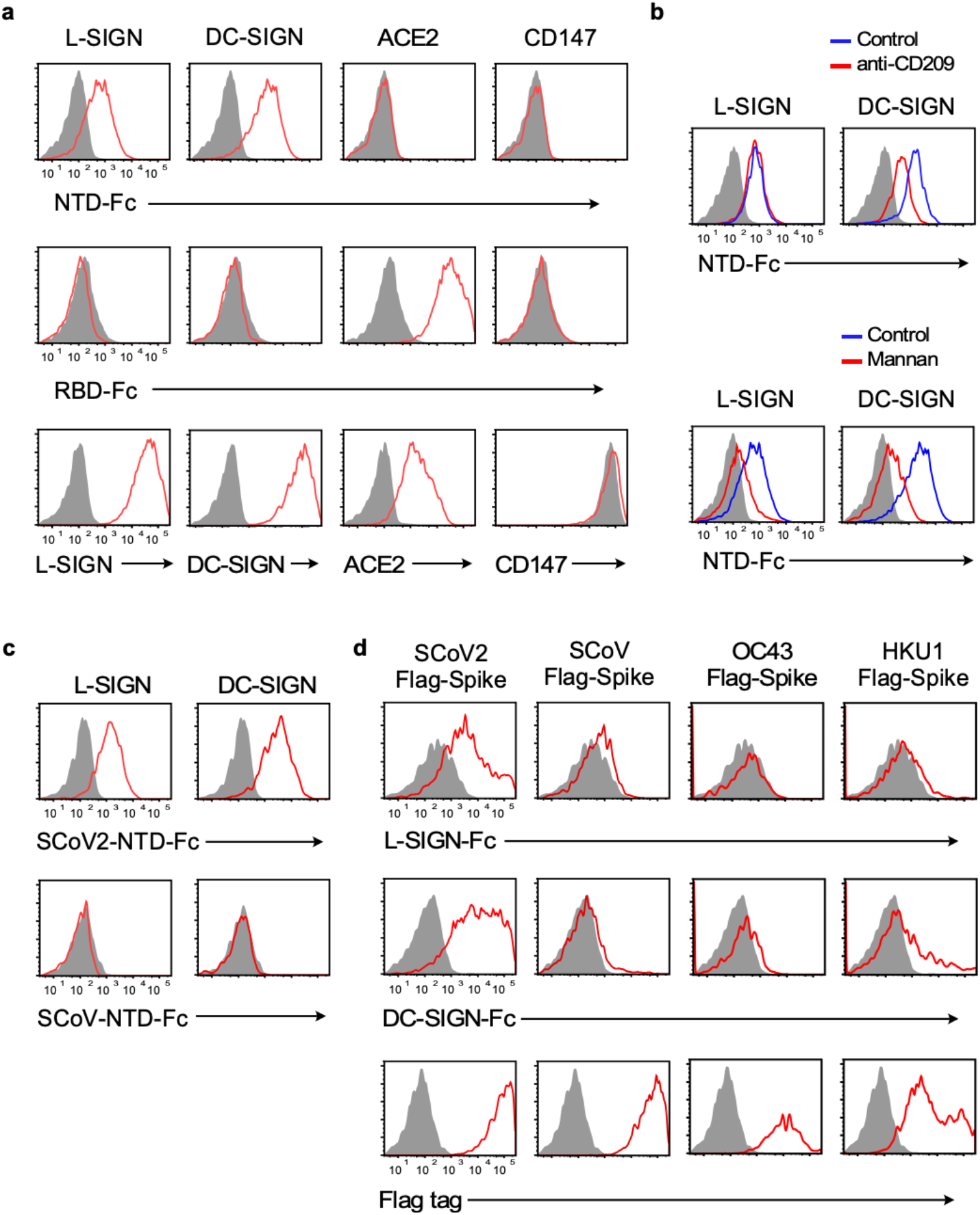
Cell surface binding analyses of the SCoV2 spike bound to DC-SIGN and L-SIGN using flow cytometry. **a,** Cells transfected with ACE2, L-SIGN, DC-SIGN, and CD147 (red line) or mock (shaded gray) were stained with SCoV2-NTD-Fc and RBD-Fc fusion protein (10 μg/mL). Each transfectant was also stained with a specific monoclonal antibody. **b,** Effect of mannan and anti-CD209 (anti-DC-SIGN) antibody on SCoV2-NTD-Fc binding to DC- and L-SIGN transfectants or mock (shaded gray). The transfectants were preincubated with mannan (light blue line) or anti-CD209 antibody (dark blue), followed by the staining with NTD-Fc fusion protein. **c,** Staining of DC- and L-SIGN transfectants (red line) or mock (shaded gray) with SCoV2-NTD-Fc and SCoV-NTD-Fc fusion proteins (10 μg/mL). **d,** Cells transfected with flag-tagged spike proteins of SCoV2, SCoV, human coronavirus OC43, or HKU1 (red line) or mock (shaded gray) were stained with DC-, L-SIGN-Fc fusion proteins or anti-Flag-tag antibody. Proportions of the stained cells are shown. Data are representative of three independent experiments.

The SCoV spike protein has a high sequence similarity (approximately 76%) to the SCoV2 spike protein. However, SCoV-NTD-Fc fusion protein did not bind to DC-SIGN or L-SIGN transfectants unlike the SCoV2-NTD-Fc fusion protein (Fig. 2c). Similarly, DC-SIGN-Fc and L-SIGN-Fc fusion proteins bound to flag-tagged SCoV2 spike transfectants, but bound weakly to SCoV spike transfectants although both transfectants were well stained with anti-flag mAb (Fig. 2d). Likewise, DC-SIGN-Fc and L-SIGN-Fc fusion proteins did not bind to the human coronaviruses, OC43 and HKU1 (Suppl. Fig. 2). These results also suggested that the NTD of the SCoV2 spike protein is specifically modified by unique glycans that mediate the binding of NTD to L-SIGN and DC-SIGN.

There are seven and eight N-glycosylation sites located on the NTD of SCoV and SCoV2, respectively (Fig. 3a). The protein sequence alignment of both NTDs revealed that there are two unique glycosylation sites, i.e., N74 and N149, on the SCoV2-NTD (Fig. 3a). Interestingly, both glycosylation sites are in the most variable region within the NTD. To identify which glycosylation sites interact with DC-SIGN, we substituted SCoV2-NTD glycosylation site Asn (N) with Gln (Q) and analyzed the binding to DC-SIGN (Fig. 3b, c). The mutation at N149Q diminished the NTD binding to DC-SIGN, while N74Q and N282Q mutation resulted in a substantial reduction of the NTD binding to DC-SIGN. These data suggest that unique glycans on N149 of NTD are responsible for the interaction with SIGNs.

**Fig. 3.**
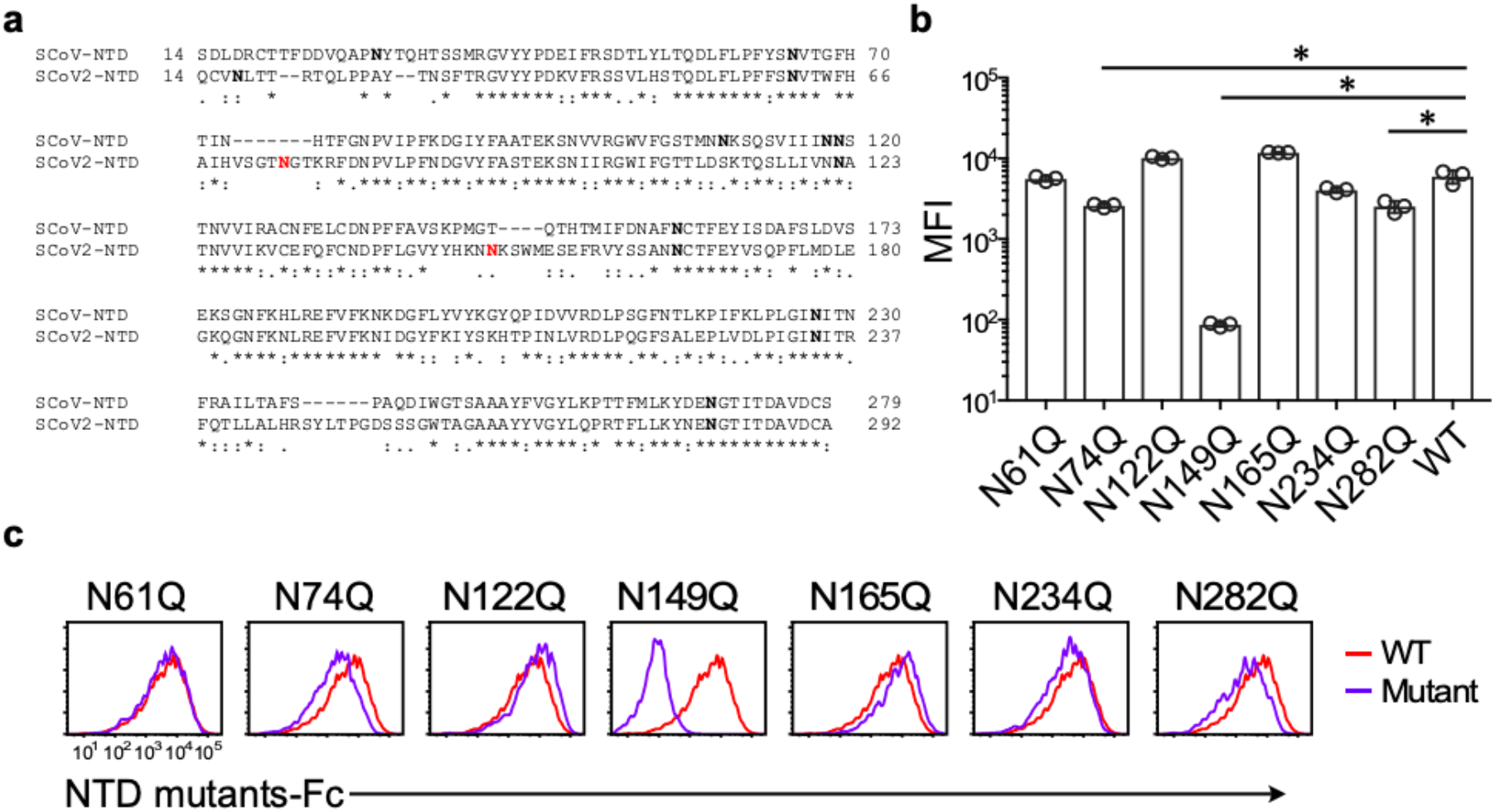
Effect of N-linked glycosylation on SCoV2-NTD interaction with DC-SIGN. **a,** Amino acid sequence alignment of the NTD between SCoV and SCoV2 spike. Bold asparagine (N) residues indicate the N-linked glycan sites ^30–32^. Highlighted asparagine residues in red indicate the unique N-glycosylation sites in SCoV2. **b,** DC-SIGN transfectants were stained with NTD-Fc fusion proteins in which N-glycosylation sites were mutated and analyzed by flow cytometry. Mean fluorescence intensities (MFI) of the cells were shown. Asterisks indicate statistical significance derived from unpaired T-test; \**P*<0.0001. **c,** Representative histogram of DC-SIGN transfectants stained with NTD-Fc mutants is shown. Proportion of cells stained with NTD-Fc mutants were shown. Data are representative of three independent experiments.

### L-SIGN and DC-SIGN mediate SCoV2 infection by inducing membrane fusion

It has been reported that L-SIGN and DC-SIGN are involved in infection by several viruses such as HIV, HCV, and SARS-CoV. To determine whether L-SIGN and DC-SIGN are involved in SCoV2 infection, we used SCoV2 spike-pseudotyped vesicular stomatitis virus (SCoV2-PV) carrying a luciferase reporter gene to infect HEK293T cells stably transfected with L-SIGN or DC-SIGN and analyzed the outcome (Fig. 4a, Suppl. Fig. 2). L-SIGN and DC-SIGN transfectants were considerably infected with SCoV2-PV. Furthermore, bona fide SCoV2 also infected L-SIGN and DC-SIGN transfectants. This indicated that DC-SIGN and L-SIGN serve as receptors for SCoV2 infection.

**Fig. 4.**
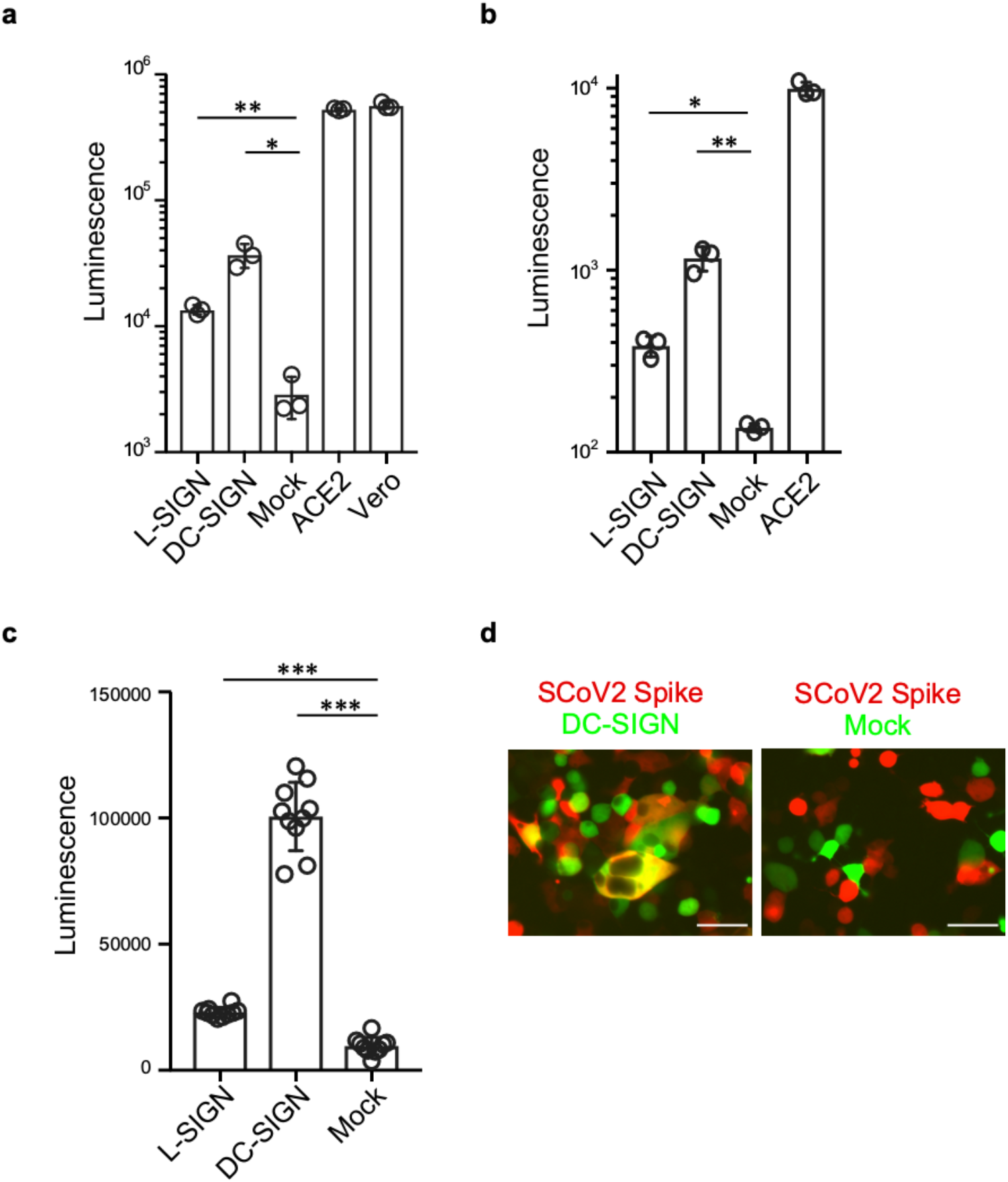
DC-SIGN and L-SIGN mediate SCoV2 infection. **a,** SCoV2-PV infection on DC-SIGN, L-SIGN, mock, or ACE2 transfectants and Vero E6 cells. SCoV2-PV carrying a luciferase gene was used for the infection and luciferase activity was measured 24 h later. Asterisks indicate statistical significance derived from unpaired T-test; \**P* = 0.0018; \*\**P* = 0.0003 **b,** DC-SIGN or L-SIGN transfectants were infected with recombinant SCoV2 with the NanoBiT luciferase gene. The luciferase activity was measured 24 h later. Asterisks indicate statistical significance derived from unpaired T-test; \**P* = 0.001; \*\**P* = 0.0006. **c,** Cell-cell fusion assay of SCoV2 spike transfectants and DC- or L-SIGN transfectants. The effector cells expressing spike and T7 polymerase were cocultured with target cells expressing DC-SIGN, L-SIGN, mock, and T7 promoter-driven luciferase. Luciferase activities were measured after 24 h. Asterisks indicate statistical significance derived from unpaired T-test \*\*\**P*<0.0001. **d,** Cell-cell fusion assay of SCoV2 spike transfectants (red) and DC-SIGN transfectants (green). Representative images are shown. Scale bar represents 50 μm in length. Data are representative of three independent experiments.

Being an enveloped virus, membrane fusion between the viral envelope of SCoV2 and the host cell membrane is a critical step in viral infection. To understand the function of L-SIGN and DC-SIGN in membrane fusion during SCoV2 infection, we used a cell-cell fusion assay using a T7 promoter-based luciferase reporter system^7,16^. In this system, we cocultured effector cells (transfected with SCoV2 spike and T7 polymerase) with target cells (transfected with DC-SIGN or L-SIGN with luciferase gene regulated by a T7 promoter) and the luciferase gene is expressed only when cell-cell fusion between effector cells and target cells is induced. Interestingly, cell-cell fusion activity occurred when effector cells were cocultured with target cells expressing L-SIGN or DC-SIGN (Fig. 4c). Additionally, when effector cells expressing SCoV2 spike protein and red fluorescence protein were cocultured with target cells expressing DC-SIGN and green fluorescence protein, fused cells expressing both red and green fluorescence proteins were detected (Fig. 4d). These data suggest that L-SIGN or DC-SIGN mediate membrane fusion during SCoV2 infection of SIGN-expressing cells.

### moDCs facilitate SCoV-2 dissemination in trans via DC-SIGN

We then sought to establish the function of endogenously expressed L-SIGN or DC-SIGN. Since L-SIGN has been reported to be expressed in type II alveolar cells, L-SIGN is likely involved in the SARS-CoV-2 infection of lungs. However, we could not find any cell lines expressing endogenous L-SIGN. On the other hand, DC-SIGN is expressed on monocyte-derived DCs (moDCs). For that reason, we isolated CD14^+^ cells from human PBMC and differentiated them into DCs in the presence of cytokines, IL-4 and GM-CSF. The moDCs expressed DC-SIGN, low CD14, high MHC-II, and CD74 (Fig. 5a). As observed with DC-SIGN transfectants (Fig. 2), SCoV2-NTD-Fc fusion protein bound to moDCs (Fig. 5b). Anti-DC-SIGN monoclonal antibodies and mannan prevented the binding of NTD-Fc to moDCs (Fig. 5c). These results indicated that DC-SIGN was expressed on primary cells associated with the NTD of the SCoV2 spike protein. However, moDCs were not infected with SCoV2-PV, but with the VSV-G PV (Fig. 5d). Recently, CD74 induced by CIITA has been reported to inhibit the membrane fusion of SCoV2^17^. The considerable expression of CD74 by moDCs (Fig. 5a) may confer resistance to SCoV2 infection. Therefore, there is likely that L-SIGN or DC-SIGN expressed in CD74 negative cells could play a critical role in SARS-CoV-2 infection.

**Fig. 5.**
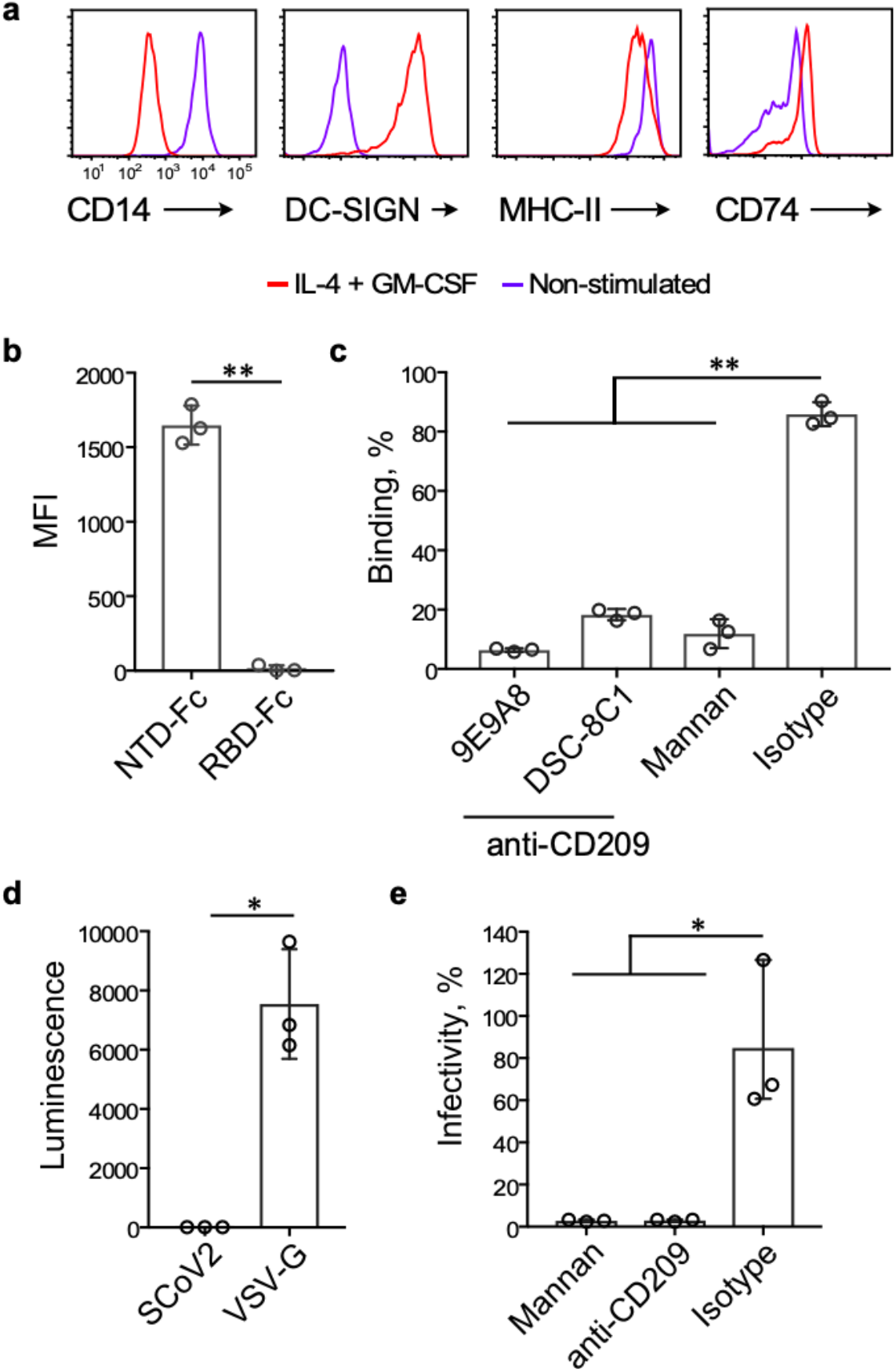
moDCs facilitate SCoV2 infection in trans through DC-SIGN. **a,** Phenotype of the CD14^+^ cells from PBMC cultured in the presence (red line) or absence (purple line) of cytokines (IL-4 and GM-CSF, 500 IU/mL each) for three days. **b,** Binding of SCoV2-NTD-Fc and RBD-Fc of SCoV2 spike to moDCs. The amount of Fc-fusion proteins bound to moDCs were presented as mean fluorescence intensity (MFI). Asterisks indicate statistical significance derived from an unpaired T-test; \*\**P*<0.0001. **c,** Blocking of SCoV2-NTD-Fc binding to moDCs by two specific anti-DC-SIGN mAbs, isotype antibody, and mannan. The percentage of binding is calculated based on the MFI value of non-blocking control. Asterisks indicate statistical significance derived from unpaired T-test; \*\**P*<0.0001. **d,** SCoV2-PV and VSV-G PV infection on moDCs. Asterisks indicate statistical significance derived from unpaired T-test; \**P* = 0.0021. **e,** SCoV2-PV infection in trans on Vero E6 cells facilitated by moDCs. moDCs was incubated with SCoV2-PV in the presence or absence of anti-DC-SIGN antibody or mannan, and cocultured with Vero E6 cells after extensive washing. Infectivity was calculated based on the luciferase activity of a non-blocking control. Asterisks indicate statistical significance derived from unpaired T-test; \**P* = 0.005. Data are representative of three independent experiments.

NTD-Fc has been observed to bind well to moDCs; therefore, SCoV2 virions bound to DC-SIGN on moDCs may be infectious to other cells. We incubated moDCs with SCoV2-PV followed by extensive washing. Thereafter, moDCs incubated with SCoV2-PV were cocultured with Vero cells. Significant infection was observed in Vero cells and the infection was completely blocked by anti-DC-SIGN antibodies or mannan (Fig. 5e). Due to the general circulation of DCs in the body, SCoV2 bound to DCs via DC-SIGN may be involved in the spread of SCoV2 throughout the body.

### Neutralization of SIGN-mediated SCoV2 infection by antibodies

Antibodies are of importance in host defense against SARS-CoV-2 infection. The function of antibodies in SIGN-mediated SARS-CoV-2 infection was evaluated. 4A8 and 4A2 are COVID-19 patient-derived anti-NTD mAbs^10^ and C144 is an anti-RBD mAb^18^ (Suppl. Fig. 3). Both 4A8 and C144 have been reported to neutralize SCoV2 infection. Surprisingly, anti-RBD C144 mAbs as well as anti-NTD 4A8 blocked SCoV2-PV infection of HEK-L-SIGN and HEK-DC-SIGN. In contrast, the 4A2 mAb did not block infection (Fig. 6a, b). These data indicate that RBD as well as NTD are involved in SCoV2 infection of SIGN-expressing cells. The effect of antibodies derived from SCoV2-infected patients was investigated; their sera contain antibodies against NTD and RBD (Suppl. Fig. 4). SCoV2-PV infection of DC-SIGN transfectants was hindered by the sera from two SCoV2-infected patients but not from healthy control serum (Fig. 6c). On the other hand, the sera could not neutralize VSV-G PV infection. (Fig. 6d). This shows that SIGN-mediated infection is counteracted by serum antibodies against spike proteins similar to ACE2-mediated infection. In addition, our data imply that antibodies against NTD as well as RBD are useful in blocking SCoV2 infection^19,20^.

**Fig. 6.**
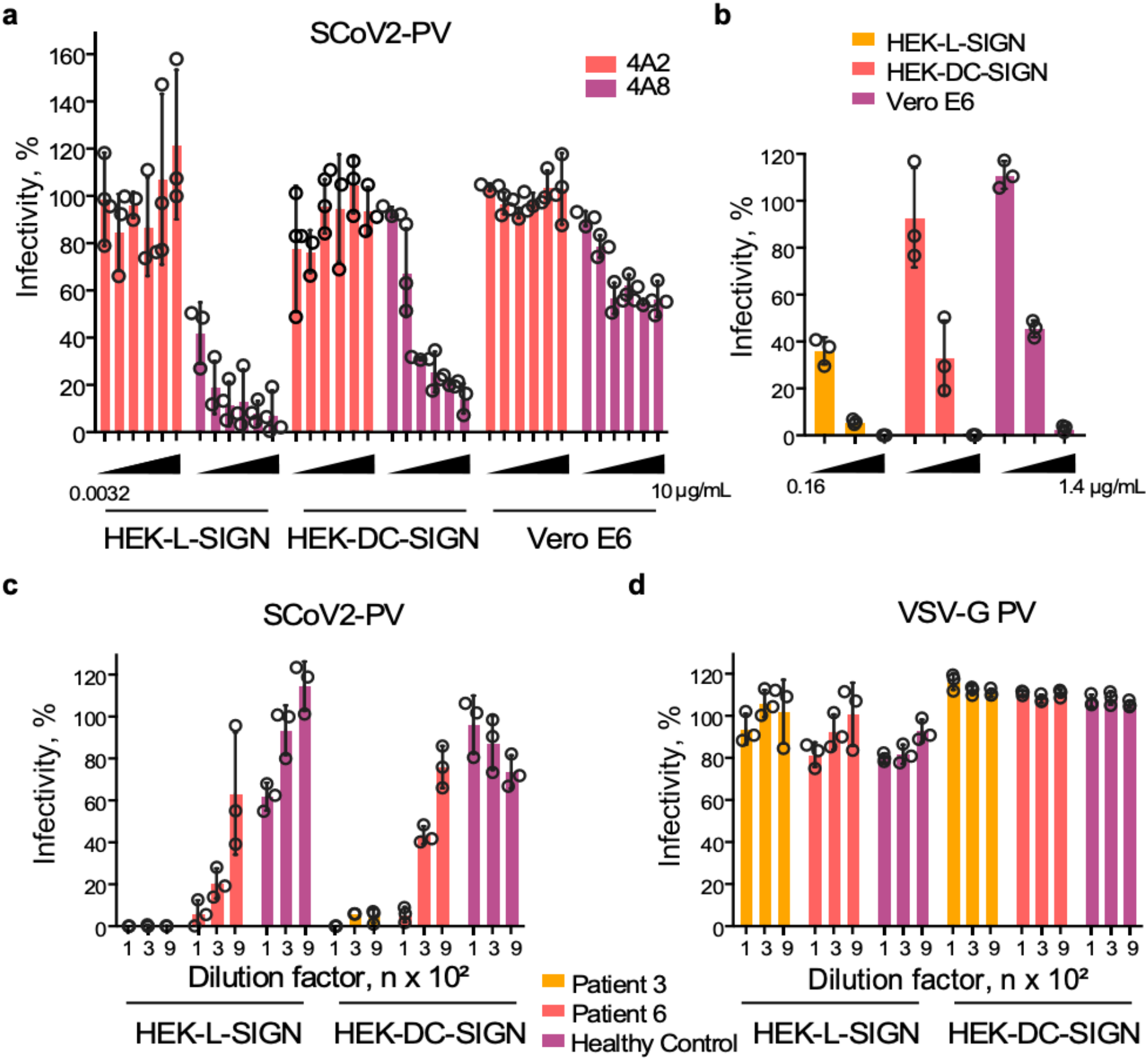
Neutralizing potency of SCoV2 convalescent patients’ sera and anti-NTD antibodies to L-SIGN and DC-SIGN mediated infection. Neutralization assay of pseudotyped SCoV2 infection on HEK-L-SIGN, HEK-DC-SIGN, and Vero cells by human anti-SCoV2-NTD mAbs **(a)** and human anti-SCoV2-RBD (C144) specific mAb **(b)**. SCoV2-PV preincubated with mAbs was used to infect the cells. Neutralization assay of SCoV2-PV **(c)** and VSV-G PV **(d)** infection of L-SIGN or DC-SIGN transfectants by serum from SCoV2-infected patients or healthy control. SCoV2-PV or VSV-G PV preincubated with sequentially diluted serum was used to infect L-SIGN or DC-SIGN transfectants and luciferase activity was measured. The percentage of infectivity was calculated based on the luminescence value of no serum or mAb control. Data are representative of three independent experiments.

## Discussion

SCoV2 is more widespread than its closest human coronavirus, SCoV^3^, and therapeutic interventions are yet to be identified. ACE2 is the primary receptor that interacts with the SCoV2 spike glycoprotein (i.e., RBD) and consequently facilitates viral entry^3^. However, the understanding of the pathogenesis of SCoV2 remains incomplete, including the mechanism of pneumonia, thrombosis, endothelitis, and cytokine storm, which are often associated with severe cases of the disease^21–23^. It is unclear how SCov2 causes pneumonia since the expression of ACE2 in the lung is relatively low and limited to type-II alveolar cells and some (<2%) of the respiratory epithelial cells^12^. Additionally, the presence of SCoV2 in non-pulmonary organs^14^ as well as the detection of viral DNA in non-ACE2 expressing cells^13^ point to an alternative receptor for SCoV2 entry and an unknown virus-spreading mechanism within the host. Recently, neuropilin has been reported to be involved in the SCoV2 cell entry by associating with the proteolytically processed spike protein. However, transfection of neuropilin alone to HEK293 cells did not mediate the SCoV2 infection, and the coexpression of ACE2 and TMPRSS2 with neuropilin was required for the SCoV2 infection^24,25^. On the other hand, L-SIGN, cloned from lung cDNA library as an NTD binding receptor, and DC-SIGN mediated SCoV2 infection in the absence of ACE2 and TMPRSS2. Indeed, L-SIGN is expressed in lung type-II alveolar cells and endothelial cells^26^. Furthermore, DCs or specialized macrophages expressing DC-SIGN can be found in the lung^27,28^. Their localization in the lung further strengthens the increasingly likely role of L-SIGN and DC-SIGN as receptors of SCoV2 and may therefore be important in the pathogenesis of pneumonia.

The interaction of the SCoV2-NTD with L-SIGN and DC-SIGN is specific and glycan dependent. Despite the NTD of SCoV being a highly glycosylated domain, our analyses indicate that the interaction between NTD and DC-SIGN or L-SIGN is apparently unique to the current SCoV2 because L-SIGN-Fc or DC-SIGN-Fc hardly bound to the spike protein of SCoV as well as human coronavirus OC43 and HKU1. Previous findings have shown that SCoV spike protein interacts with DC-SIGN and L-SIGN albeit with a low binding efficiency^26,29^. Similarly, we observed a very weak binding of DC- and L-SIGN-Fc to SCoV spike protein, whereas the binding to the SCoV2 spike was stronger. The overall glycosylation profile of the SCoV2 spike protein is quite different from that of SCoV^30–32^ with the N-linked glycan at residue 149 being the major glycan involved in the DC-SIGN and NTD interaction. The N-linked glycosylation site at residue 149 of SCoV2-NTD is absent in SCoV-NTD, although other N-linked glycosylation sites are conserved in SCoV-NTD. This may be the reason why L- and DC-SIGN interactions with NTD are more apparent in the current SCoV2. Indeed, both L-SIGN and DC-SIGN recognize a pentasaccharide motif (GlcNAc2Man3)^33^ that is also present in the glycans at N149 of SCoV2-NTD^30^.

We demonstrated that L-SIGN and DC-SIGN mediate both the pseudotyped and bona fide SARS-CoV-2 infections. Furthermore, both L-SIGN and DC-SIGN induce cell-cell fusion via their association with the spike protein. SCoV2 requires proteolytic cleavage of the spike protein by the host proteases (e.g., TMPRSS2) after binding to an entry receptor^3^. Besides, the lysosomal protease, cathepsin L was shown to proteolytically activate the SCoV spike protein during viral entry thereby facilitating membrane fusion with the host cell membrane^34^. Recent research revealed that the thyroglobulin domain (cathepsin L inhibitor) of the CD74 p41 isoform blocks the cathepsin L-mediated viral entry of SCoV2^17^. CD74 is an essential molecule for the proper assembling of MHC class II molecules and its expression is regulated by CIITA, a principal transcription factor for MHC class II expression^35^. Because moDCs express both CD74 and MHC class II molecules favorably, CD74 appears to be responsible for the resistance of moDCs to SCoV2 infection. However, expression of CD74 is largely limited to immune cells such as B cells, monocytes, and dendritic cells, while most cells including endothelial cells do not express CD74. Therefore, L-SIGN or DC-SIGN expressing cells that lack CD74 expression appear to be susceptible to SCoV2 infection.

We also show that moDCs facilitate pseudotyped SCoV2 infection in Vero cells in trans. Trans-infection by DCs is well reported in HIV infection^36^. Due to their migration in the body, DCs may function as transporters of SCoV2 by associating with the virus via DC-SIGN to infect cells expressing ACE2 or L-SIGN, which in turn facilitates virus dissemination in the host. This might explain the presence of viral protein in other organs, including the liver^14^, where L-SIGN is highly expressed in human liver sinusoidal endothelial cells that do not express CD74^37^. Furthermore, infection of the endothelial cells of blood vessels may consequently lead to thrombosis, which is frequently linked to severe disease conditions^22^. Several human tissues express similar mRNA expression levels of L-SIGN as those of the lung, making them likely targets of SCoV2 infection.

The presence of neutralizing antibodies for DC-SIGN-mediated SCoV2 infection in COVID-19 patients implies the relevance of NTD as a therapeutic target besides RBD. Notably, the majority of the isolated mAb from SCoV2 patients did not recognize RBD and those mAbs that neutralize SCoV2 failed to prevent binding of the spike to ACE2^10^. In our study, the NTD-specific mAb, 4A8, neutralized SIGN-mediated infection of SCoV2, suggesting the therapeutic potential of anti-NTD antibodies against SIGN-mediated infection. More importantly, anti-RBD mAb also inhibited the SCoV2-PV infection of DC-SIGN and L-SIGN expressing cells. Although HEK293 cells do not express ACE2, certain molecules on HEK293 cells may interact with RBD becoming involved in infection of SIGN-expressing cells. This may suggest that L-SIGN or DC-SIGN is required for the infection of ACE2-low or ACE2-negative cells, where levels are not enough to mediate SCoV2 infection. Further analysis of L-SIGN or DC-SIGN-mediated infection would be important to understand the etiology of SARS-CoV-2-related diseases. Also, targeting SIGN-mediated infection as well as ACE2-mediated infection would be important for effective vaccine development.

## Methods

### Antibodies

Mouse anti-CD209 mAb (clone 9E9A8), mouse IgG2a isotype control antibody (BioLegend, San Diego, CA, USA), mouse anti-L-SIGN mAb (clone 19F7), mouse anti-MHCII (clone WR18) (Santa Cruz Biotechnology, Dallas, TX, USA), mouse anti-human ACE2 mAb (R&D Systems, Minneapolis, MN, USA), rat anti-Flag mAb (Sigma-Aldrich, St Louis, MI, USA), mouse anti-human CD14-APC mAb (eBioscience, San Diego, CA, USA), Donkey anti-mouse IgG-APC mAb, anti-human IgG-Fc fragment specific-APC mAb, anti-rat IgG-APC mAb (Jackson ImmunoResearch, West Grove, PA, USA).

### Cell lines and cell culture

The following cell lines were used in this study: Vero E6 is an African Green Monkey Kidney cell line, HEK293T cells are a human kidney cell line, HEK293T-DC-SIGN is a stable transfectant of HEK293T expressing DC-SIGN, HEK293T-L-SIGN is a stable transfectant of HEK293T expressing L-SIGN, HEK293T-ACE2 is a stable transfectant of HEK293T expressing ACE2, HEK293T-L-SIGN/TMPRSS2 is a stable transfectant of HEK293T expressing L-SIGN and TMPRSS2, HEK293T-DC-SIGN/TMPRSS2 is a stable transfectant of HEK293T expressing L-SIGN and TMPRSS2. Stable cell lines were prepared using retrovirus transfection produced from Plat-E^38^. Briefly, Plat-E cells were co-transfected with target gene vectors and an amphotropic envelope vector using polyethylenimine (PEI; Polysciences, Warrington, PA, USA) as described previously^39^. Culture supernatant containing retroviruses was harvested after 48 hours and premixed with DOTAP liposomal transfection reagent (Roche, Basel, Switzerland) prior to spin transfection at 2400 rpm at 32°C for 2 hours into target cells. The transfected cells were cultured for three days and subsequently sorted using an SH800S cell sorter (Sony, Minato City, Tokyo, Japan). All cell lines were maintained in DMEM (Nacalai Tesque, Kyoto, Japan) supplemented with 10% heat-inactivated fetal bovine serum (FBS) (Biological Industries, Cromwell, CT, USA), penicillin (100 U/mL), and streptomycin (100 μg/mL) (Nacalai Tesque, Japan) and cultured at 37°C in 5% CO_2_.

### Genes

The followings genes were used in the current study, SARS-CoV-2 spike (amino acid from 1– 1273, GenBank accession NC_045512.2), SARS-CoV spike (amino acid from 1–1255, GenBank accession AY278491), human coronavirus HKU1 spike (amino acid from 1–1352, GenBank accession NC_006213), human coronavirus OC43 (amino acid from 1–1353, GenBank accession KY674921) DC-SIGN (GenBank accession NM_021155.4), L-SIGN (gene obtained from lung cDNA library screening corresponding to GenBank accession NM_001144904.2), human ACE2 (GenBank accession NM_021804.3), human TMPRSS2 (GenBank accession BC051839.1), human CD147 (GenBank accession NG_007468), human IgG1 Fc segment^6,40,41^, genes for variable regions of mAbs, C144, 4A2, and 4A8, were retrieved from previous studies^10,18^.

### cDNA library screening

A retrovirus cDNA library was constructed based on a previous study^42^. Briefly, cDNA was generated from a human lung poly-A^+^ RNA (cat. 636105, Takara Bio, Kusatsu, Shiga, Japan) using a cDNA library construction kit (Takara Bio, Japan) and cloned into EcoRI and NotI sites of pMXs retrovirus vector. The ligated cDNA was transformed into XL10-gold and the plasmids were purified using a plasmid plus midi kit (Qiagen, Germantown, MD, USA). The complexity of the cDNA library was 2 × 10^6^. The cDNA library was then transfected into PLAT-E packaging cells to generate a retrovirus cDNA library. Subsequently, the retrovirus cDNA library was transfected into 2B4 mouse T cell hybridoma and sorted for cells binding to the SCoV2-NTD-Fc fusion protein using an SH800S cell sorter (Sony, Japan). Single-cell clones were obtained and the genomic DNA was then isolated using a Wizard SV genomic DNA purification kit (Promega, Madison, WI, USA). The target gene was amplified by PCR using pMXs vector forward (5’GGTGGACCATCCTCTAGA3’) and reverse (5’CCCTTTTTCTGGAGACTAAAT3’) primers. The identity of the target gene was determined by DNA sequencing. The L-SIGN gene obtained corresponded to GenBank accession NM_001144904.2.

### Mutagenesis of N-linked glycosylation sites of SCoV2-NTD

Single point SCoV2-NTD-Fc N-linked glycosylation site mutants (N61, 74, 122, 149, 165, 234, 282Q) were generated using a QuikChange site-directed mutagenesis kit (Agilent Technologies, Santa Clara, CA, USA) to change the respective asparagine (N, Asn) residue to glutamine (Q, Gln).

### Recombinant protein production

Recombinant DC-SIGN-Fc, L-SIGN-Fc, SCoV-NTD-Fc (amino acids 14–279), SCoV2-NTD-Fc (amino acids 14–292), SCoV2-RBD-Fc (amino acids 335–587), and SCoV2-NTD-Fc N-glycosylation site mutant fusion proteins were independently expressed as a secreted protein using HEK293T cells. The recombinant mAbs, C144, 4A2, and 4A8 were produced using Expi293 cells. The Fc fusion protein in the culture supernatant was quantified using a protein A beads assay. Essentially, the protein A was coupled to 4 μm aldehyde/sulfate latex beads (Invitrogen, USA) according to the manufacturer’s instructions and used to capture the Fc fusion protein in the culture supernatant. The captured Fc fusion proteins were then detected using anti-human IgG-Fc conjugated to APC and measured using flow cytometry. The protein concentration was calculated using a known concentration of IgG-Fc fusion protein as a standard. For recombinant proteins that required purification, the culture supernatant containing the secreted fusion proteins was purified using protein-A conjugated Sepharose beads (Cytiva, Marlborough, MA, USA).

### Flow cytometry analysis

FACSVerse™ or FACSCalibur flow cytometer (Becton Dickinson, Franklin Lake, NJ, USA) was used for flow cytometry analysis. For the binding study, cells were transiently co-transfected with the plasmid containing the respective receptors and GFP using a PEI transfection reagent^39^. Forty-eight hours post-infection, cells were then stained with Fc fusion proteins for 30 minutes at 4°C in Hank’s buffer containing 0.1% BSA and identified using goat anti-human IgG-Fc-APC Ab prior to flow cytometry analysis. For DC- and L-SIGN blocking experiments, the cells were pre-blocked with 5 μg/mL of anti-CD209 mAb or 50 μg/mL of mannan (Sigma, USA) for 30 minutes before staining with Fc fusion protein. For the staining of the spike protein, DC-SIGN-Fc (5 μg/mL) or L-SIGN-Fc fusion protein (20 μg/mL) premixed with goat anti-human IgG-Fc-APC mAb (3 μg/mL) was used. For the analysis of SCoV2-NTD N-linked glycosylation site mutants’ interaction with DC-SIGN, 10 μg/mL of each mutant was used. Surface-expressed SCoV2-NTD (amino acids 14–333), SCoV2-RBD (amino acids 335–587), and SCoV2-S2 (amino acids 588– 1219) were generated by fusing the respective domains with a transmembrane region derived from PILRα at the C-terminal^6^.

### Cell-cell fusion assay

Effector cells were prepared by transiently transfecting T7 RNA polymerase plasmid, pCAG-T7, and SCoV2 spike plasmid into HEK293T cells. For the target cells, HEK293T cells were transiently co-transfected with T7 promoter-luciferase plasmid, pT7EMCV-Luc, and DC-SIGN in a pcDNA3.4 vector, L-SIGN in a pcDNA3.4 vector, or a pcDNA3.4 empty vector (mock). After two days of transfection, the effector cells (1 × 10^4^ cells) were cocultured with the target cells (1 × 10^4^ cells) in a 384-well white plate (Greiner Bio-One, Kremsmünster, Austria). The cells were then incubated for 24 hours at 37°C in 5% CO_2_ and the luciferase activity was measured using the ONE-Glo™ luciferase assay (Promega, USA) according to the manufacturer’s instructions. The signals were recorded using a luminescence plate reader TriStar LB941 (Berthold Technologies, Bad Wilbad, Germany). For the cell-cell fusion assay using fluorescence protein, effector cells were prepared by transiently co-transfecting with SCoV2 spike and red fluorescence protein (pCI-neo-DsREDExp) into HEK293T cells. For target cells, HEK293T cells were transiently co-transfected with DC-SIGN and green fluorescence protein (pMX-GFP). After two days of transfection, the effector and target cells were enriched using a Sony SH800 cell sorter (Sony, Japan) prior to co-culture as described above. Cells were then analyzed using an IX83 Olympus microscope (Olympus, Shinjuku City, Tokyo, Japan).

### SCoV2 spike-pseudotyped virus

HEK293T cells were transiently transfected with expression plasmids for the SCoV2 spike protein. Twenty-four hours post-transfection, VSV-G-deficient vesicular stomatitis virus (VSV) carrying a luciferase reporter gene complemented in trans with VSV-G protein^43^ was added and incubated for two hours. Cells were then carefully washed with DMEM media without FBS and incubation continued with DMEM supplemented with FBS at 37°C in 5% CO_2_ for 48 hours. The supernatant containing the pseudotyped SCoV2 virions was harvested and aliquoted before storage at −80°C.

### CD14^+^ cell preparation and moDCs differentiation

Peripheral blood mononuclear cells (PBMCs) were prepared from healthy individuals by centrifugation using Ficoll-Paque™ PLUS (GE Healthcare, Chicago, IL, USA). CD14^+^ cells were isolated by positive selection using anti-CD14 conjugated microbeads (Miltenyi Biotec, Bergisch Gladbach, Germany) according to the manufacturer’s instructions. Monocyte-derived dendritic cells (moDCs) were stimulated as previously described^44^. Briefly, 5 × 10^5^ cells/mL of CD14^+^ enriched cells were stimulated using recombinant human (rh) GM-CSF and rhIL-4 (BioLegend, USA) at a concentration of each 500 IU/mL in RPMI1640 media supplemented with 10% FCS at 37°C in 5% CO_2_. The expression of CD209 was detected after two days of stimulation using an anti-CD209 monoclonal antibody.

### Direct infection assay

For pseudotyped SCoV2 infection, 5 × 10^3^ cells were mixed with the pseudovirus for 24 hours at 37°C in 5% CO_2_ in a 384-well plate. Luciferase activity was then measured using the ONE-Glo™ luciferase assay (Promega, USA) according to the manufacturer’s instructions. The signals were recorded using a luminescence plate reader TriStar LB941 (Berthold Technologies, Germany). For neutralization assay, essentially the same protocol was conducted with the exception that the pseudovirus was preincubated with diluted patient sera or anti-NTD, clone 4A8 and 4A2 monoclonal antibodies for 30 minutes at room temperature. SARS-CoV-2 convalescent patients’ sera were collected from Osaka University Hospital. Patients’ sera collection was approved by the IRB (No. 19546). Recombinant SCoV2 virus with a high-affinity NanoBiT (HiBiT) luciferase gene in the N-terminus of the ORF6 gene was produced as previously described^45^. The virus (MOI 3) was added to 1.5 × 10^4^ cells for 24 hours at 37°C, 5% CO_2_ in a 96-well plate. SCoV2 infected cells were collected and luciferase activity was then measured using a Nano-Glo HiBiT lytic assay system (Promega, USA), according to the manufacturer’s instructions. The signals were recorded using a luminometer.

### Trans-infection assay

moDCs were incubated with pseudotyped SCoV2 at 37°C for 60 minutes. For the blocking experiment, moDCs were preincubated with anti-CD209 antibodies (5 μg/mL), mouse IgG2a antibodies (Isotype control), or mannan (100 μg/mL) before incubation with pseudovirus. After incubation, the cells were washed three times with DMEM and subsequently added to Vero cells at a 1:1 ratio. The cells were incubated at 37°C in 5% CO_2_ for 24 hours and luciferase activity was measured using a ONE-Glo™ luciferase assay (Promega, USA) according to the manufacturer’s instructions.

### Sequence, data, and statistical analysis

The amino acid multiple sequence alignment in Fig. 3a was generated using the CLUSTAL Omega multiple sequence alignment tool from NCBI. FlowJo (BD Biosciences, San Jose, CA, USA) was used for analyzing flow cytometry data and Graphpad Prism version 7.0e was used for graph generation and statistical analysis. Asterisks indicate statistical significance.

## Acknowledgements

Soh W.T. is supported under the Kishimoto Foundation Fellowship. We would like to thank Dr. Masako KOHYAMA and Dr. Wataru NAKAI, Akihito SAKOGUCHI for technical advice. We also thank Akemi ARAKAWA, Asa TADA, Sumiko MATSUOKA for their excellent technical supports. This work was supported by JSPS KAAKENHI Grant Numbers JP18H05279, JP19H03478, MEXT KAKENHI Grant Numbers JP19H04808, Japan Agency for Medical Research and Development (AMED) under Grant Number 19fk0108161h0001, 20nf0101623h0201, 20nk0101602h0201.

## Author contributions

W.T.S., M.Y., S.T., and H.A. conceived experiments. W.T.S., Y.L., N.E.E., and O.C., performed experiments. N. H. collected patients’ sera. W.T.S. and H.A. wrote the manuscript. All authors read, edit, and approved the manuscript.

